# Human hippocampal theta oscillations reflect sequential dependencies during spatial planning

**DOI:** 10.1101/372011

**Authors:** Raphael Kaplan, Adrià Tauste Campo, Daniel Bush, John King, Alessandro Principe, Raphael Koster, Miguel Ley-Nacher, Rodrigo Rocamora, Karl J. Friston

## Abstract

Movement-related theta oscillations in rodent hippocampus coordinate ‘forward sweeps’ of location-specific neural activity that could be used to evaluate spatial trajectories online. This raises the possibility that increases in human hippocampal theta power accompany the evaluation of upcoming spatial choices. To test this hypothesis, we measured neural oscillations during a spatial planning task that closely resembles a perceptual decision-making paradigm. In this task, participants searched visually for the shortest path between a start and goal location in novel mazes that contained multiple choice points, and were subsequently asked to make a spatial decision at one of those choice points. We observed ~4-8 Hz hippocampal/medial temporal lobe theta power increases specific to sequential planning that were negatively correlated with subsequent decision speed, where decision speed was inversely correlated with choice accuracy. These results implicate the hippocampal theta rhythm in decision tree search during planning in novel environments.

## Introduction

Recent evidence has linked the hippocampus with planning in rodents (Miller et al., 2017) and humans (Kaplan et al., 2017a). Moreover, changes in hippocampal theta power (approx. 4-8Hz in humans) have been observed during memory-guided decision-making in well-learned environments in both species (Guitart-Masip et al., 2013; Schmidt et al., 2013; Belchior et al., 2014). However, it remains unclear whether changes in hippocampal theta power are associated with planning in novel environments. Notably, rodent type I movement-related hippocampal theta oscillations (Vanderwolf, 1969) are linked to sweeps of place cell activity produced by hippocampal theta phase precession (O’Keefe & Recce, 1993). It has been hypothesized that these ‘theta sweeps’ could serve as a mechanism to plan trajectories online (Johnson & Redish, 2007; Wikenheiser & Redish, 2015; Watrous et al., 2018). This raises the possibility that similar increases in human hippocampal theta power are induced by the planning of forward trajectories.

To investigate the role of the hippocampal theta rhythm in online spatial planning (i.e., the search of decision trees), we created a spatial task that required little to no learning, in which participants could draw upon their experience in the physical world (Kaplan et al., 2017a). We tested human participants on this task while recording from the hippocampus either invasively, using intracranial electroencephalography (iEEG); or non-invasively, using whole-head magnetoencephalography (MEG). In both cases, participants were instructed to search for the shortest path between a start and goal in novel mazes that afforded multiple paths. Participants were then asked which direction they would take from one of two choice points along the shortest path (Fig. 1).

**Fig 1.**
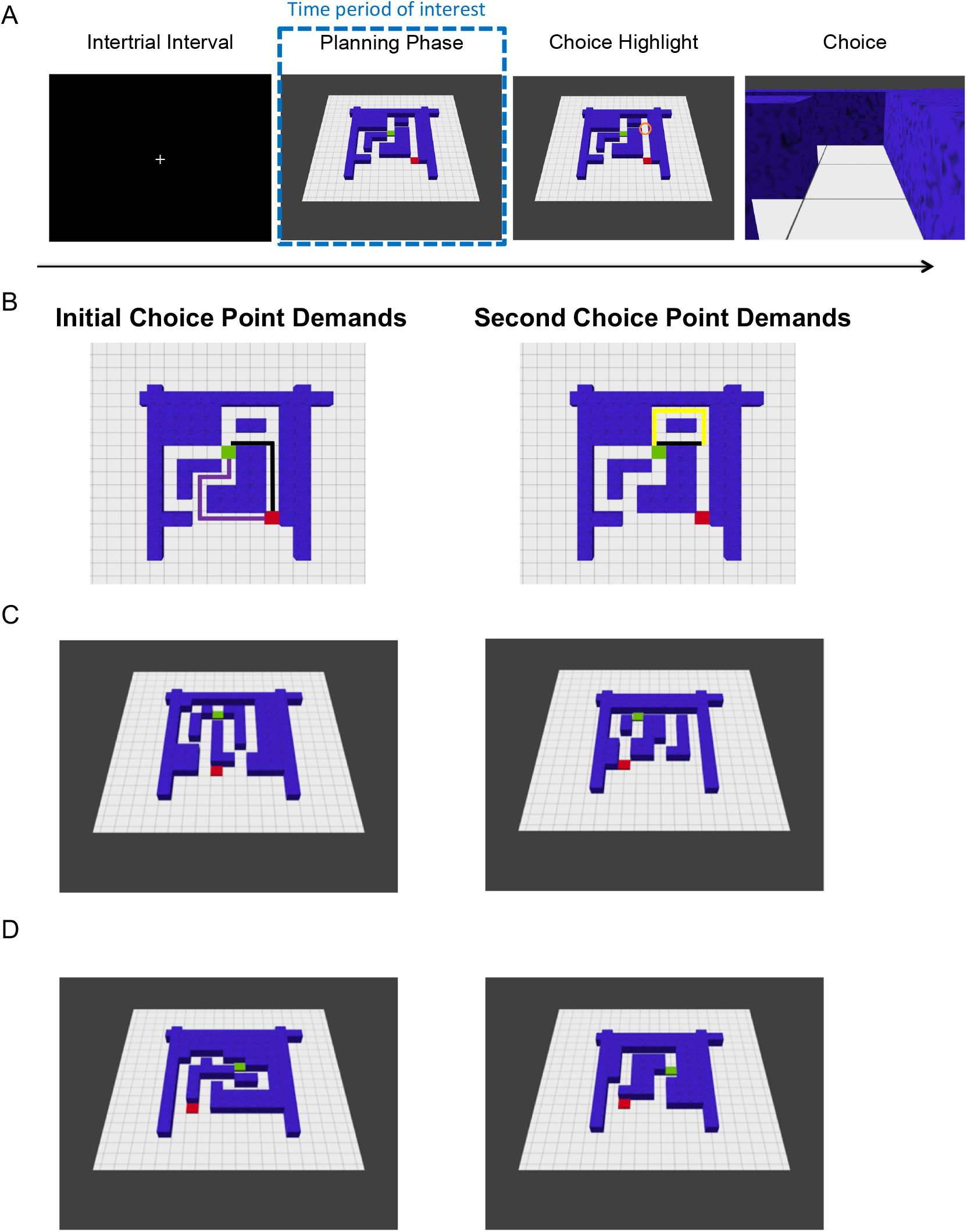
Task. A. Each trial (i.e., visually presented maze) began with an inter-trial interval (ITI) of 1.5s. Next, during a 3.25s planning phase, participants had to infer the shortest path from a start point (red square) to a goal location (green square) and remember the chosen direction for each choice point along the shortest path. A choice point was subsequently highlighted (choice highlight) for 250ms. This was either the first (i.e. initial) or second (i.e. subsequent) choice point along the shortest path. Participants were then asked which direction (e.g., left or forward) they would take at that choice point during a choice period that was cued by a first-person viewpoint of the highlighted location. Participants had a maximum of 1.5 s to make their choice using a button box. B. Overhead view (not shown during the experiment) of the maze in A, indicating which path lengths contribute to initial and second choice point demands (black line represents shortest path). C. Left: Example sequential planning trial with a small path length difference (demanding) at the red square/initial choice point and large (less demanding) path length difference at the second choice point. Right: Example trial with a large (less demanding) path length difference at the red square/initial choice point and small (demanding) path length difference at the second choice point. D. Left: Example non-sequential (control) trial with a small path length difference (demanding). Right: Example non-sequential (control) trial with a large path length difference (less demanding).

Crucially, the mazes were designed to induce forward planning in terms of a two-level tree search, where participants needed to maintain the decisions they made at each choice point. At both choice points, there was a small, medium, or large path length difference – creating a total of (3×3) nine conditions allowing us to test the effect of planning demands at each choice point depth (i.e., initial or second). In parallel, our task also contained a non-sequential control condition, where participants were presented with mazes containing only one choice point (Fig. 1D). In either case, we associate a smaller path difference with greater ambiguity and processing demands. Importantly, in any trial, participants were only prompted to make one choice after seeing the full maze; however, until the choice point was highlighted, they did not know which decision (i.e. either the initial or second/subsequent choice point along the correct path for sequential mazes) would be probed in sequential planning trials (Fig. 1). After planning their route, participants were asked to choose—at a specified choice point—the direction of the shortest path to the goal location (Fig. 1). This provided a measure (reaction time, RT) with which to quantify their (subjective) uncertainty to complement the (objective) difference in path lengths. This design allowed us to ask whether hippocampal theta power is selectively related to demands at specific choice points and how the theta rhythm relates to successful sequential spatial planning.

## Results

### Behavioral Performance

Twenty-two participants in the MEG study made correct choices on 87.9 ± 6.13% of sequential planning trials (mean ± SD; control trials: 86.4 ± 4.95%), with an average reaction time (RT) of 469 ± 99ms (control trials: 363 ± 112ms). Paired t-tests showed that their RT was significantly higher for sequential than non-sequential (i.e. control) trials (t(21)=9.55; p<.001), without any difference in accuracy (t(21)=1.42; p=.171). In addition, RTs were strongly inversely correlated with accuracy across MEG participants in both sequential (t(21)=-5.72; p<0.001) and non-sequential control trials (t(21)=-5.72; p<.001). After accounting for planning demands induced by the path length differences at each choice point (mean path length differences at the two choice points), RTs were still negatively correlated with accuracy in both sequential (t(21)=-5.25; p<.001) and non-sequential control trials (t(21)=-5.14; p<.001). In other words, participants responded faster when they made accurate choices. Moreover, these results demonstrate that RTs directly relate to accurate performance on the spatial planning task.

We then asked whether accuracy and RT were specifically influenced by path length differences and choice point depth, with the aim of disentangling the effects of first/initial versus second/subsequent choice point demands on planning accuracy and RT. Using a repeated measures ANOVA, we looked for an effect of path length difference and choice point depth on accuracy and RTs in MEG participants. We observed a main effect of path length on both accuracy (F(2,20)=9.09; p=.002; Fig. 2A) and RTs (F(2,20)=5.06;p=.017; Fig. 2B), driven by higher accuracy and faster RTs for larger path length differences; as well as a significant interaction between initial (i.e. first) and second (i.e. subsequent) choice points and path length differences on both accuracy (F(4,18)=11.0; p<0.001) and RTs (F(4,18)=4.75; p=0.009). Post-hoc t-tests revealed that this interaction resulted from medium path length differences being significantly less demanding (i.e. producing higher accuracy and faster RTs) when they were at the initial, as opposed to the second, choice point (Accuracy: t(21)=3.62; p=.002; RT: t(21)=-4.17; p<.001).

**Figure 2:**
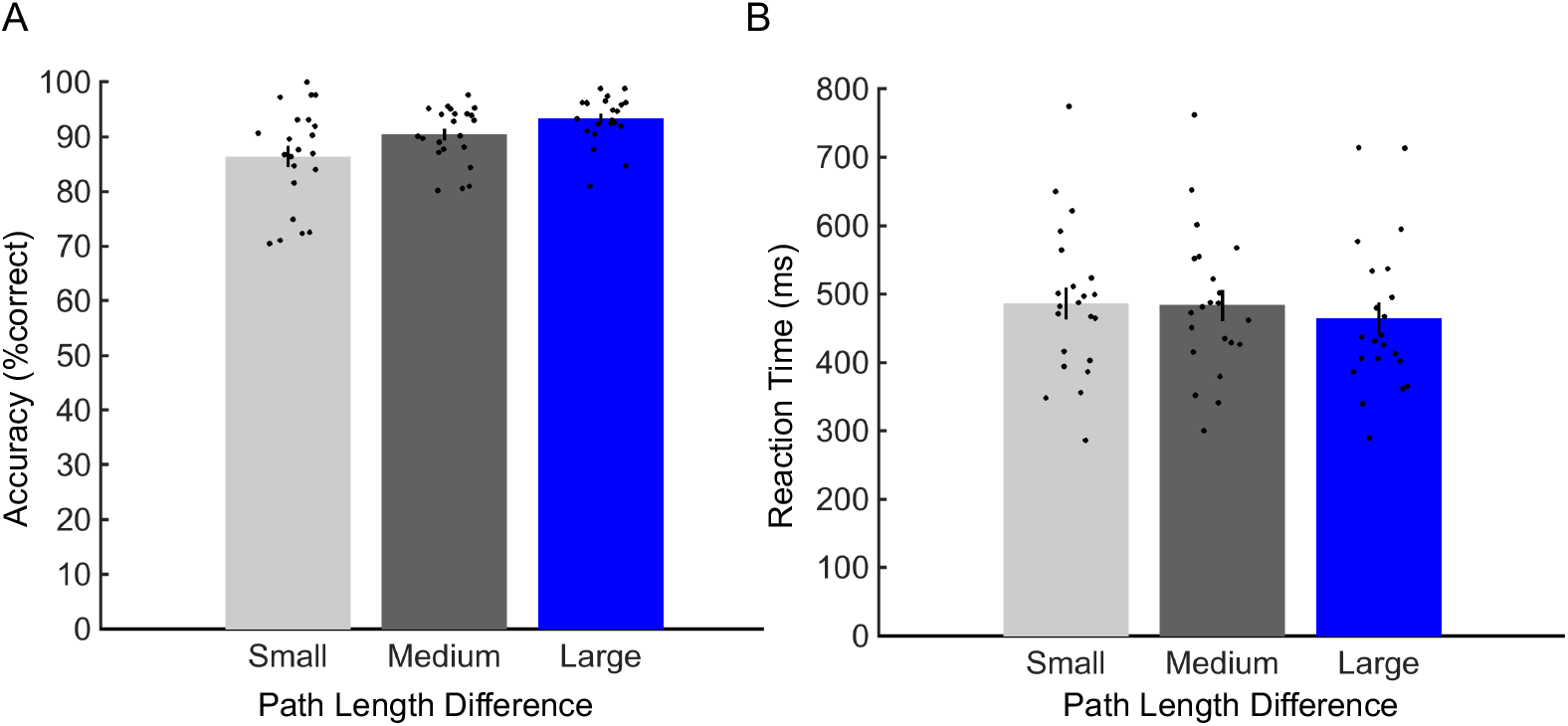
Behavior. A. Accuracy. Left: Significant main effect (p=0.002) of path length differences (small, medium, and large) on choice accuracy, collapsed across first and second choice points. B. Reaction time. Significant main effect (p=0.017) of path length differences (small, medium, and large) on reaction times, collapsed across initial and second choice point. All error bars show ± SEM.

### MEG Analyses

Using MEG source reconstruction, we asked whether power changes in five canonical frequency bands (delta / low theta: 1-3 Hz, theta: 4-8Hz, alpha: 9-12Hz, beta: 13-30Hz, and gamma: 30-80Hz) anywhere in the brain were related to differences in spatial planning. Focusing on RTs, we found a significant negative correlation between 4-8Hz theta power during the sequential planning phase and subsequent RTs in a left hippocampal cluster (x:-36, y:-20, z:-20, t(21)=-4.28; small volume corrected (SVC) peak-voxel p=.011; Fig. 3A). Specifically, increased hippocampal theta power during planning periods preceded faster decisions – an effect that was also observable at the scalp level (Fig. 3C). Notably, we did not observe any correlation between theta power and trial-by-trial choice accuracy anywhere in the brain, although this is likely due to a relatively small number of errors.

**Fig 3.**
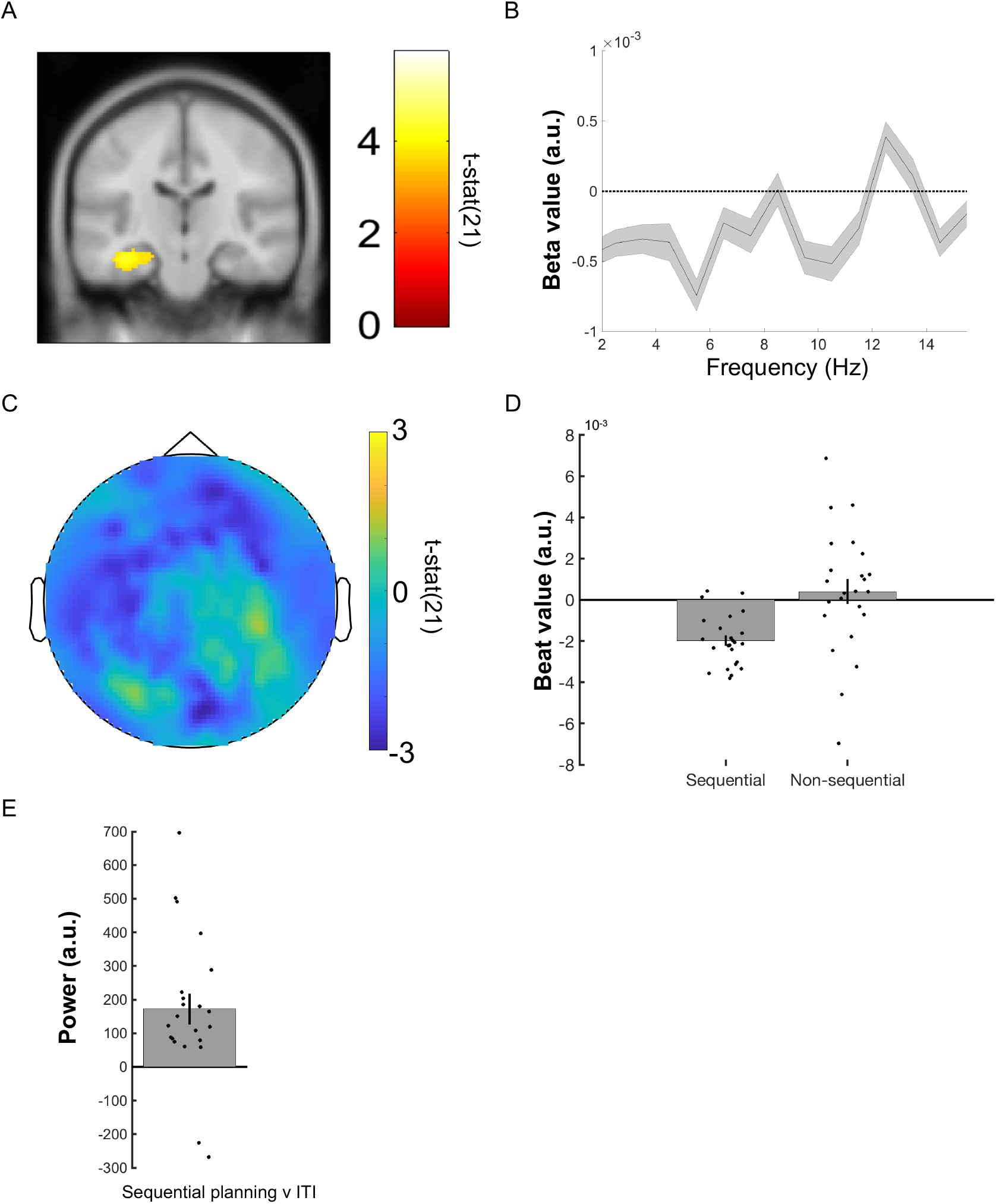
Reaction time correlation with MEG theta power. A. Linearly Constrained Minimum Variance (LCMV) beamformer source reconstruction image showing significant 4-8 Hz left hippocampal theta power source negative correlation with RT (x:-36, y:-20, z:-20) in 22 healthy participants. Images displayed at the statistical threshold of p<0.001 uncorrected for visualization purposes. B. Beta value spectrum from 1 to 15 Hz for hippocampal RT theta power effect showing peak negative correlation in the 4-8 Hz theta band. C. Negative 4-8 Hz theta power correlation with RT shown at the scalp level for 22 healthy participants. D. Data from a 10 mm sphere around left hippocampal peak voxel from RT contrast for both sequential and non-sequential/control planning trials. E. Data from a 10 mm sphere around left hippocampal peak voxel from RT contrast showing increased theta power during planning phase versus the ITI period. All error bars show ± SEM.

In addition, we found a significant negative correlation between theta power and RTs in the right ventral temporal lobe (x:36, y:-42, z:-26; t(21)=4.49; family wise error (FWE) corrected peak-voxel p=.012; Fig. S1), which extended into posterior parahippocampal cortex. We did not observe a significant positive correlation between 4-8Hz planning period theta power and subsequent RTs anywhere in the brain. Elsewhere, we observed 9-12Hz alpha power changes in the right occipital lobe/cerebellum that negatively correlated with RT (x:28, y:-70, z:-22; t(21)=-5.99; FWE corrected peak-voxel p=.014; Fig. S1). However, we observed no other significant correlations between oscillatory power and RT in any other brain regions or frequency band.

To assay whether significant power changes related specifically to sequential planning, we tested whether each correlation described above was stronger for sequential planning trials versus non-sequential/control trials. Using a 10mm sphere around the respective peak voxels, we observed that hippocampal RT theta effects selectively corresponded to sequential planning (t(21)=-2.33; p=.03; Fig. 3D), while right ventral temporal/parahippocampal theta (t(21)=-1.38; p=.181; Fig. S1) and occipital/cerebellar alpha effects did not (t(21)=-1.74; p=.095; Fig. S1).

We then asked whether sequential spatial planning was associated with a general increase in left hippocampal theta power. Again, using a 10mm sphere around the left hippocampal peak, we observed a significant increase in 4-8Hz hippocampal theta power in this region during the sequential planning period versus ITI (t(21)=3.74; p=.001; Fig. 3E). Conducting the same sequential planning versus ITI analysis in the other areas exhibiting RT effects, we observed significant increases in both ventral temporal lobe theta (t(21)=2.79; p=.011) and occipital alpha (t(21)=4.44; p<.001) power during sequential planning.

Finally, isolating hippocampal theta power changes, we tested for the effects of processing demands (path length differences) at initial and second/subsequent choice points (e.g., quicker RT for mazes with less demanding initial choice points). Using a repeated measures ANOVA (path length difference by choice point depth), we tested whether the left hippocampal region (exhibiting a theta power correlation with RT) also showed an effect of path length differences at initial versus second choice points related to RT. We did not observe any significant effect of path length difference by choice point depth in the left hippocampus (F(4,18)=1.79; p=.175), or any other brain region.

### Hippocampal depth recordings

Next, to verify our source reconstructed MEG effects, we examined changes in low frequency oscillatory power during the 3.25s sequential planning period using iEEG recordings from hippocampal depth electrodes (Fig. 4A) of a single high performing pre-surgical epilepsy patient (95.5% accuracy; mean RT: 423 ± 123ms). Notably, a hippocampal theta rhythm was readily visible in the raw iEEG traces during this planning phase (Fig. 4B). Further validating our MEG results, we asked whether iEEG hippocampal theta power during sequential planning correlated with the patient’s subsequent RT. Interestingly, we observed a negative correlation between ~3-6 Hz hippocampal theta power during the 3.25s planning phase and subsequent RT (r=-0.194; p=.043; Fig. 4C), although this result should be interpreted with caution given the relatively small number of measurements. Overall, we observed hippocampal oscillations during the sequential planning period that were most prominent in the theta band and exhibited power increases in the same frequency band that correlated with faster subsequent RT in the MEG dataset.

**Fig 4.**
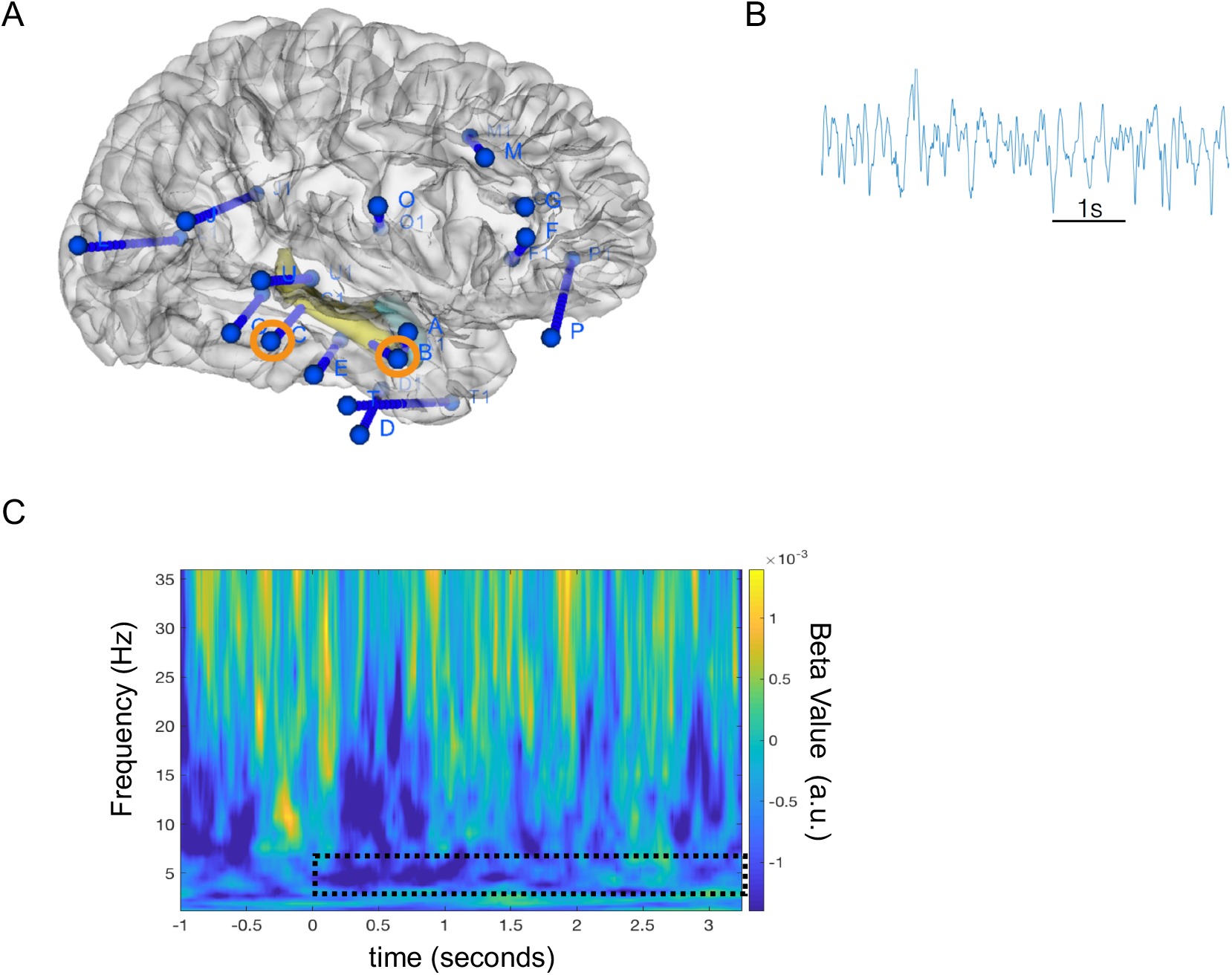
Intracranial EEG data from hippocampal depth electrodes. A. Image of electrode locations in the patient overlaid on 3D brain template. Right hippocampal depth electrodes with contacts used in the present analyses are highlighted in orange. B. Sample raw trace showing prominent ongoing theta band oscillations during the spatial planning task. C. Time-frequency plot showing a negative correlation over trials between subsequent reaction time (RT) and 3-6 Hz theta power during sequential planning period (highlighted with dotted black box) averaged across both hippocampal contacts.

## Discussion

We examined how the human hippocampal theta rhythm relates to planning sequential decisions in novel environments. Linking hippocampal theta to participants’ performance on a spatial planning task, theta power during the planning phase correlated with faster subsequent spatial decisions in MEG and iEEG participants (Figs. 3 & 4). Furthermore, decision speed correlated with choice accuracy, regardless of path length differences. Linking the human hippocampal theta rhythm to processing demands, we found that hippocampal theta power selectively corresponded to planning performance in mazes containing multiple choice points during the MEG task. Here, we relate our findings to the extant hippocampal decision-making literature and speculate on potential computational roles associated with the hippocampal/medial temporal lobe theta rhythm.

Our observation of increased hippocampal theta power during spatial decision-making adds to an emerging literature investigating the role of the hippocampal theta rhythm during decision-making in rodents (Johnson & Redish, 2007; Schmidt et al., 2013; Belchior et al., 2014; Wikenheiser & Redish, 2015; Pezzulo et al., 2017) and humans (Guitart-Masip et al., 2013). Yet, the specific role of the hippocampal theta rhythm in planning has remained unclear; despite recent evidence relating the rodent (Miller et al., 2017) and human hippocampus (Kaplan et al., 2017a) to planning. Additional support for a hippocampal role in planning comes from evidence that hippocampal neurons code the distance to goal locations (Ekstrom et al., 2003; Villette et al., 2015; Sarel et al., 2017; Watrous et al., 2018). Furthermore, Wikenheiser and Redish (2015) found that firing of place cell sequences coupled to the hippocampal theta rhythm extended further on journeys to distal goal locations. We parallel these findings by showing that hippocampal theta power was selectively related to efficient sequential planning, which further implicates the human hippocampal rhythm in prospective evaluation of upcoming choices during planning.

Differing from previous MEG/iEEG hippocampal theta studies that observe power increases related generally to enhanced task performance (Lega et al., 2012; Olsen et al., 2013; Backus et al., 2016; Heusser et al., 2016), we find hippocampal theta power effects associated with planning behavior in sequential, but not simpler mazes. Given the known relationship between the hippocampal theta rhythm and spatial trajectories, these findings may relate to sequential spatial decision-making that focuses on signifying a ‘location’ update within a sequence of choices. Supporting this explanation, recent work has suggested that the hippocampus can suppress noise in our everyday environment to focus on sub-goals during multi-step planning (Botvinick & Weinstein, 2014). Furthermore, biophysical models predict that the hippocampal theta rhythm can underlie this type of ‘sub-goaling’ within deep/sequential planning by updating our location from initial starting points to subsequent sub-goals (Kaplan & Friston, 2018).

Still, several aspects of our results remain unclear. For instance, an alternative explanation for not observing right hemisphere or non-sequential hippocampal theta power spatial planning effects could be that there are multiple theta sources corresponding to sequential and non-sequential RT effects (Miller et al., 2018), which MEG does not have adequate spatial resolution to resolve. Work comparing potential hemispheric or anterior/posterior differences in the hippocampal theta rhythm may help address this question (Miller et al., 2018). Furthermore, the direct relationship between behaviorally relevant hippocampal theta power changes and the reactivation of place cell sequences is not well characterized, since we are not measuring single-neuron activity. However, Watrous and colleagues (2018) recently observed that human hippocampal single units exhibit phase-locking to the theta rhythm and that this phase-locking encoded information about goal locations during virtual navigation. Work building on this line of research –using hippocampal iEEG recordings to inform whole-brain non-invasive MEG analyses – could provide a novel way to potentially answer questions about the role of the hippocampal theta rhythm in spatial decision-making.

Our task is reminiscent of perceptual decision-making paradigms and there is an emerging link between saccadic searches and the hippocampus. However, it should be noted that we only measured electrooculogram (EOG) signals during this task, not saccadic behavior. Future work can build on the growing literature linking visual exploration to movement-initiated hippocampal activity (MacIver et al., 2017). Of particular interest, Wang and colleagues (2018) found that the firing of single neurons in the human MTL related to successful visual searches for a target item embedded within an image. Moreover, recent studies of neural oscillations in the hippocampal formation in humans and non-human primates have related saccadic exploration of visual space to spatial exploration of the physical world (Jutras et al., 2013; Staudigl et al., 2018). Yet, how these findings relate to sequential decision-making/planning remains unclear.

We studied multi-step planning in an explicitly spatial domain, but it isn’t known whether updating our ‘location’ to subsequent choice points relates more to the overhead visual searches of the maze or a more abstract decision space (Schiller et al., 2015; Kaplan et al., 2017b). On one hand, there is mounting evidence of the type I movement-related rodent hippocampal theta rhythm extending to virtual (Ekstrom et al., 2003, 2005; Watrous et al., 2011; Kaplan et al., 2012; Bush et al, 2017; Watrous et al., 2018) and real-life navigation in humans (Aghajan et al., 2017; Bohbot et al., 2017). However, evidence from non-spatial domains is lacking. Potential clues may come from the investigation of the role of the hippocampal formation in imagined exploration of spatial environments (Byrne et al., 2007; Bellmund et al., 2016; Horner et al., 2016; Kaplan et al., 2017c). Indeed, the hippocampal theta rhythm has been observed during teleportation from one location to another (Vass et al., 2016) – providing further support for a role of the hippocampal theta rhythm in navigating more abstract spaces. Future work exploring the role of the hippocampal theta rhythm in both perceptual exploration (Jutras et al., 2013; Aronov et al., 2017) and prospective evaluation during abstract sequential decisions (Kaplan et al., 2017b), can determine how generalizable spatial navigation-related hippocampal theta effects are to other abstract spaces (Lisman & Redish, 2009).

In summary, our findings suggest that the human hippocampal theta rhythm plays an important role during spatial decision-making in novel environments. Namely, our data relate hippocampal theta power changes to sequential dependencies during spatial planning. Moreover, we present findings from a spatial decision-making task that more closely resembles perceptual decision-making than virtual navigation paradigms. This therefore leaves open the possibility that the human hippocampal theta rhythm also relates to prospective evaluation during multi-step decisions in non-spatial domains.

## Supplemental Information

### Supplemental Experimental Procedures

#### Participants

##### MEG

Twenty-four participants (14 female: mean age 23.5 yrs; SD of 3.49 years) gave written consent and were compensated for performing the experimental task, as approved by the local research ethics committee at University College London in accordance with Declaration of Helsinki protocols. All participants had normal or corrected-to-normal vision and reported to be in good health with no prior history of neurological disease. Due to technical difficulties, two participants were removed from our sample, leaving twenty-two participants in the behavioral and MEG analyses presented here.

##### iEEG

Pre-surgical EEG recordings from 2 patients with pharmacoresistant focal-onset seizures and hippocampal depth electrodes gave written consent, as approved by the local ethics committee at Hospital del Mar and in accordance with Declaration of Helsinki protocols. One patient was removed from analyses, because of visual difficulties due to an inferior occipital lesion, leaving one patient with normal vision presented in the current analysis. A summary of the patient’s characteristics is given in Table 1.

**Table 1.** Patient information

| Age/<br>Sex | Handedness | Seizure<br>Onset/Freq | Education | Epileptic<br>Focus | Drugs & Dosage | First-<br>language |
| --- | --- | --- | --- | --- | --- | --- |
| 23M | R (but used<br>L due to IV) | 16 yo (1<br>seizure per<br>week and<br>now seizure<br>free) | Secondary | R<br>Temporobasal<br>(temporal<br>pole) | Eslicarbazepinole<br>1000 mg/per day;<br>leviteracetam<br>1500/2x day;<br>Perampanel 8<br>mg/per day | Spanish |

All diagnostic and surgical procedures were approved by the clinical ethics committee of Hospital del Mar in accordance with the principles expressed by the Declaration of Helsinki. Electrode locations were determined solely by clinical criteria, ascertained by visual inspection of post-implantation MRI scans using Slicer 4 (Fedorov et al., 2012; www.slicer.org) and verified by an fMRI expert (R.Ka.). Patients were seizure free for at least 24 h before participation and underwent an extensive neuropsychological evaluation to check for any cognitive impairments.

#### Experimental Design

Stimuli were presented via a digital LCD projector on a screen (height, 32 cm; width, 42 cm; distance from participant, ~70 cm) inside a magnetically shielded room using the Cogent (http://www.vislab.ucl.ac.uk/cogent.php) toolbox running in MATLAB (Mathworks, Natick, MA, USA). Over the course of 220 trials, participants viewed 220 different mazes from a slightly tilted (overhead) viewpoint and later chose from first-person viewpoints within mazes generated using Blender (http://www.blender.org). All mazes had a starting location (a red square) towards the bottom of the maze and a goal location (a green square) further into the maze (Kaplan et al., 2017a). Mazes differed by hierarchical depth (number of paths to a goal location): there were 110 mazes with four possible routes (sequential/deep mazes) and a further 110 non-sequential control mazes with two possible routes (shallow mazes).

In the scanner, participants were first presented with pictures of novel mazes (Fig. 1) of varying difficulty (from an overhead viewpoint) and then asked to determine the shortest path from a starting location (a red square) at the bottom of the screen to the goal location (a green square). The overhead view appeared on the screen for 3.25 s, after which a location (choice point) along the path was highlighted briefly for 250 ms with an orange circle. The choice point location could either be the starting location or a second (subsequent) choice point. Crucially, participants would only have to make a decision about one choice point for each trial.

At either choice point, it was necessary to choose between two possible directions, which could be left, forward, or right, with an additional option to select equal, if both routes were the same distance. No second choice points with two incorrect choices were ever highlighted, only a second choice point along the optimal path after the starting location could be highlighted (Kaplan et al., 2017a). After the choice point was highlighted, a “zoomed in” viewpoint of this location (always one square back and facing the same direction as the overhead viewpoint) was presented. Depending on the possible direction at the location, participants had less than 1,500 ms (2,000 ms for the iEEG patient) to decide whether to go left, forward, right, or occasionally either direction. If no button press was made within the allotted duration, the trial counted as an incorrect trial and the experiment moved on to the 1500-ms inter-trial interval (ITI) phase. Participants never received any feedback or reward for making the correct choice. As soon as participants chose a direction, the ITI phase of a trial began. Participants repeated this trial sequence 110 times per session, for a total of two sessions. Sessions lasted approximately 10–15 min. Session order was counterbalanced between participants.

All participants completed a brief practice session consisting of 40 mazes/trials before the experiment (on a laptop outside of the scanner). Sequential mazes contained two branch/choice points between routes further in the maze, and the path length to reach the two choice points further in the maze was always equal. Mazes had square tiled floors and were 8 x 8, 9 x 9, or 10 x 10 squares in total area. In sequential mazes, we used a 3×3 factorial design. Path length differences were split between 2 (small difference), 4 (medium difference), or 6 (large difference) squares (for an example, see square tiles in the mazes presented in Fig 1) for the two paths at the starting location and a path length difference of 2, 4, or 6 squares at the optimal choice point in the maze. There was one catch trial for deep/sequential and shallow/control mazes in each session, each containing all equal path lengths (path length differences of 0). In sum, sequential maze trials could be 2, 2; 2, 4; 2, 6; 4, 2; 4, 4; 4, 6; 6, 2; 6, 4; 6, 6; (e.g. 4, 2 would have a medium path length difference of 4 at the starting location, whereas the second choice point would have a small path length difference of 2). Half of the trials in the experiment were control/shallow mazes, which only contained one choice point at the red starting square. For these mazes, path length differences were split between 2, 4, and 6, with one catch trial per session having equal path lengths.

#### iEEG recordings and artifact detection

All recordings were performed using a standard clinical EEG system (XLTEK, subsidiary of Natus Medical, Pleasanton, CA) with a 500 Hz sampling rate. A unilateral implantation was performed accordingly, using 15 intracerebral electrodes (Dixi Médical, Besançon, France; diameter: 0.8 mm; 5 to 15 contacts, 2 mm long, 1.5 mm apart) that were stereotactically inserted using robotic guidance (ROSA, Medtech Surgical, New York, NY).

Intracranial EEG signals were processed in the referential recording configuration (i.e., each signal was referred to a common reference; Tauste Campo et al., 2018). All recordings were subjected to a zero phase, 400th order finite impulse response (FIR) band-pass filter to focus on our frequency range of interest (0.5-48 Hz) and remove the effect of alternating current (Bush et al., 2017). Audio triggers produced by the stimulus presentation laptop were recorded on the monitoring system, allowing the EEG to be aligned with task information sampled at 25 Hz.

Analysis of EEG recordings focused on the 3.25 s planning periods with an additional 1.5 s baseline prior to trial onset (ITI period). All trials that included interictal spikes (IIS) or other artifacts, either within the period of interest or during the padding windows, were excluded from all analyses presented here. A 500 ms padding window was used at either end of planning period time series to minimize edge effects in subsequent analyses.

#### iEEG Time-Frequency Analysis

Estimates of dynamic oscillatory power during periods of interest were obtained by convolving the EEG signal with a seven-cycle Morlet wavelet and squaring the absolute value of the convolved signal. The wavelet transform was preferred to the Fourier transform as it does not assume stationarity in EEG recordings. To generate power spectra, the mean of dynamic oscillatory power estimates was taken over the time window of interest in the deepest contact in each hippocampal electrode. To perform baseline correction on time–frequency data for display purposes, power values were averaged across ITI periods for each frequency band, and those average values were subtracted from the power values at each time point in the planning period (Bush et al., 2017). To examine changes in oscillatory power within specific frequency bands and assess correlations among oscillatory power in each trial with RT, dynamic estimates of oscillatory power were calculated over the time and frequency windows of interest. Power values were then averaged across both hippocampal contacts to provide a single value at each time and frequency point for the patient.

#### MEG recording and preprocessing

Data were recorded continuously from 274 axial gradiometers using a CTF Omega whole-head system at a sampling rate of 600 Hz in third-order gradient configuration. Participants were also fitted with four electroculogram (EOG) electrodes to measure vertical and horizontal eye movements. MEG data analyses made use of custom made Matlab scripts, SPM8 & 12 (Wellcome Centre for Human Neuroimaging, London), and Fieldtrip (Litvak et al., 2011; Oostenveld et al., 2011). For preprocessing, MEG data was epoched into 2s baseline periods prior to the planning phase for each of the nine sequential planning conditions of interest and the three non-sequential planning control conditions. Trials were visually inspected, with any trial featuring head movement or muscular artefacts being removed.

#### MEG Source Reconstruction

The linearly constrained minimum variance (LCMV) scalar beamformer spatial filter algorithm was used to generate source activity maps in a 10-mm grid (Barnes et al., 2003). Coregistration to MNI coordinates was based on nasion, left and right preauricular fiducial points. The forward model was derived from a single-shell model (Nolte, 2003) fit to the inner skull surface of the inverse normalized SPM template. The beamformer source reconstruction algorithm consists of two stages: first, based on the data covariance and lead field structure, weights are calculated which linearly map sensor data to each source location; and second, a summary statistic based on the mean oscillatory power between experimental conditions is calculated for each voxel. Due to the proximity of frontal and anterior medial temporal lobe regions to the eyes, we wished to control for any possible influence of EOG muscular artefacts during the planning phase on estimates of oscillatory power. We therefore computed the variance of two simultaneously recorded EOG signals across each planning phase and removed any covariance between these EOG variance values and oscillatory power measurements across voxels by linear regression (Kaplan et al., 2014, 2017c). This left ‘residual’ oscillatory power measurements for all trials whose variance could not be accounted for by changes in the EOG signal between trials, and these residual values were used as summary images for subsequent analyses. RT was included as an additional nuisance regressor for the theta power source analysis investigating the effect of path length differences at different choice points. Including RT as a nuisance regressor specifically for this analysis helped determine whether there were any residual hippocampal theta power effects related to choice point demands during the planning period.

#### Statistical Analyses

There were two main periods of interest, the 1.5s ITI and 3.25s planning phase. For each of the 9 sequential planning regressors of interest (i.e., maze with a small, medium, or large path length at the second and first choice points), there were parametric regressors based on RT and accuracy (whether the choice was correctly answered; 1=incorrect choice; 2=correct choice). Inferences about these effects were based upon t- and F-tests using the standard summary statistic approach for second level random effects analysis.

A peak voxel significance threshold of p<0.05 FWE corrected for multiple comparisons was used for MEG source analyses. Given the previously hypothesized role of the hippocampus theta rhythm in planning, we report whether peak-voxels in these regions survive small-volume correction for multiple comparisons (*p* < 0.05) based on a bilateral ROI of the hippocampus (mask created using Neurosynth, Yarkoni et al., 2011). All images are displayed at the p<0.001 uncorrected threshold for illustrative purposes. Additionally, only clusters containing a significant peak voxel are displayed.

Post hoc statistical analyses were conducted using 10-mm radius spheres around the respective peak voxel specified in the GLM analysis. This allowed us to compare the effects of different regressors of interest (e.g., to determine whether a second choice point demand effect was present in a region defined by an orthogonal main effect of RT). This ensured we did not make any biased inferences in our post hoc analyses.

## Supplemental Figures

**Fig S1.**
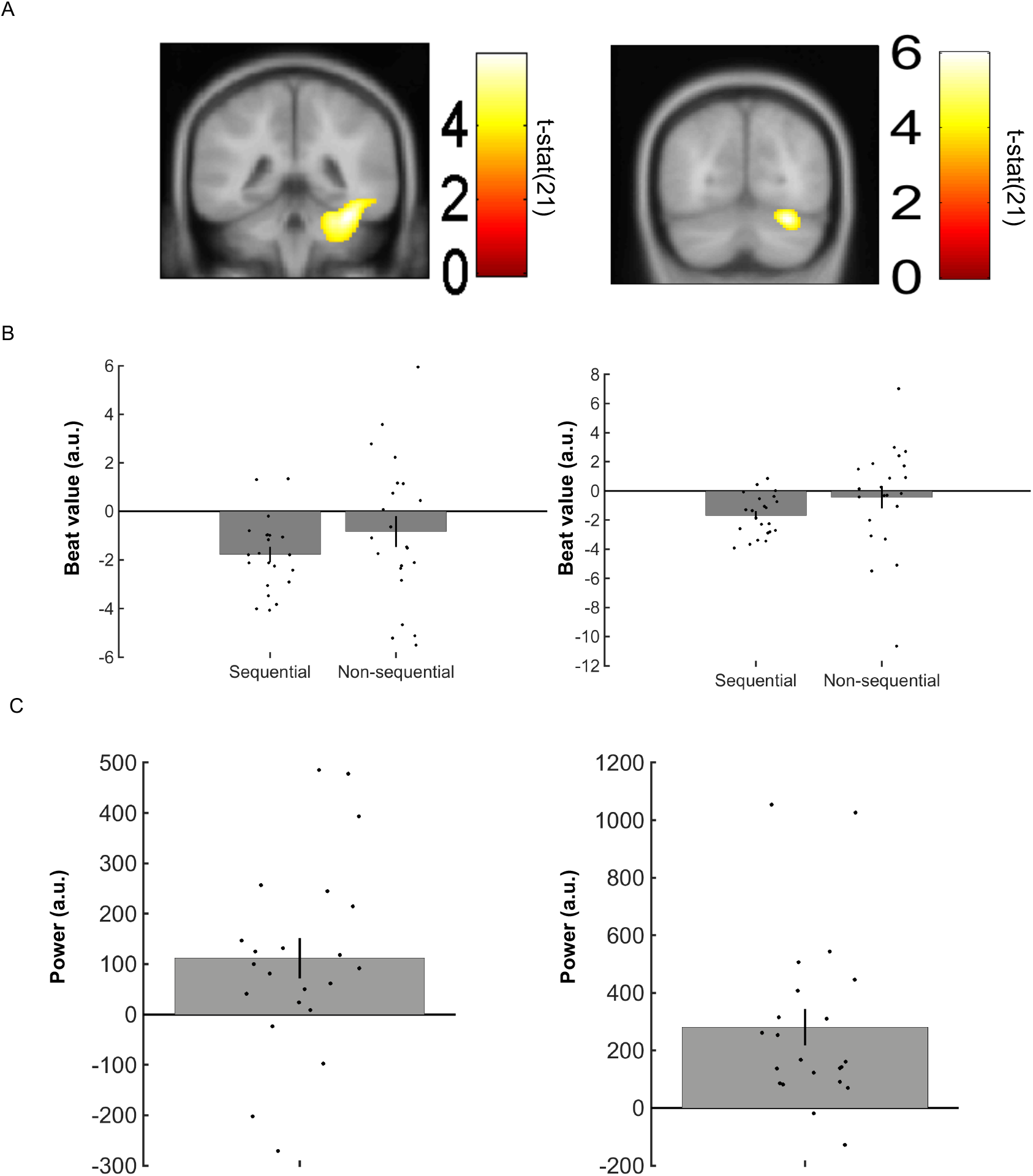
Additional reaction time correlations with MEG theta and alpha power. A. Linearly Constrained Minimum Variance (LCMV) beamformer source reconstruction images. Left: Shows significant 4-8 Hz right ventral temporal cortex theta power source negative correlation with RT (x:36, y:-42, z:-26) in 22 healthy participants. Right: Shows significant 9-12 Hz right occipital/cerebellar cortex alpha power source negative correlation with RT (x:28, y:-70, z:-22). Images displayed at the threshold of p<0.001 uncorrected for visualization purposes. B. Left: Data from a 10 mm sphere around right ventral temporal peak voxel from RT contrast for both sequential and non-sequential/control planning trials. Right: Data from a 10 mm sphere around right occipital peak voxel from RT contrast for both sequential and non-sequential/control planning trials. C. Left: Data from a 10 mm sphere around right ventral temporal peak voxel from RT contrast showing increased theta power during planning phase versus the ITI period. Right: Data from a 10 mm sphere around right occipital peak voxel from RT contrast showing increased theta power during planning phase versus the ITI period. All error bars show ± SEM.

## Acknowledgements

The research was supported by a Sir Henry Wellcome Postdoctoral Fellowship to RKa (Ref: 101261/Z/13/Z) and a Wellcome Principal Research Fellowship to KJF (Ref: 088130/Z/09/Z). We thank Carmen Pérez Enríquez for helpful discussion and the staff at Hospital del Mar for help with patients. We would also like to thank David Bradbury and Letty Manyande for assistance with MEG experimental setup. We also thank the Wellcome Centre for Human Neuroimaging for providing facilities.

